# Storing and analyzing a genome on a blockchain

**DOI:** 10.1101/2020.03.03.975334

**Authors:** Gamze Gürsoy, Charlotte M Brannon, Sarah Wagner, Mark Gerstein

**Author notes:** These authors contributed equally to this study (Gamze Gürsoy, Charlotte M Brannon).

## Abstract

The genomic characterization of individuals promises to be immensely useful for biomedical research and healthcare. However, a critical barrier to expanding personal genome sequencing is achieving secure, high-integrity storage of raw data. While cloud storage offers solutions to access such data from any place and device, the vulnerabilities of centralized storage in relation to security, data integrity, and robustness, such as single points of failure, have not yet been addressed. Blockchain is a potential alternative to these storage modes. However, storing large-scale data on blockchain can be challenging due to slow transaction speeds, the potential for chains to reach large sizes, and limitations on querying data stored on-chain. Currently, several genomic storage applications incorporate blockchain, but likely because of these challenges, many use blockchain only to facilitate and log data-access transactions, rather than to store raw genomic data on-chain. While this secures the process of data access, it does not secure the data itself, which is often stored off-chain (i.e. in a cloud or file-hosting services). Here, we developed a novel method of storing reference-aligned reads on-chain in a private blockchain network. We also developed tools for accessing and analyzing the on-chain data. We addressed the challenges of on-chain data storage by minimizing the data inserted to the chain using reference-based data compression techniques and by binning the on-chain data by genomic location to reduce retrieval times. Our tools provide open-source blockchain-based storage and access for advanced genomic analyses such as variant calling.

## Introduction

Modern advances in personalized medicine have resulted in an increasing number of individuals willing to sequence their own genome for disease-risk predictions and ancestry analysis, which has brought us closer to an era of genomic data-driven health care and biomedical research. Given the widespread interest in understanding one’s own genomic data, and the promise of these data for advancing biomedical research, it is almost inevitable that genome sequencing will become part of routine clinical care in the future and that the number of sequenced human genomes will continue to grow (Khan and Mittleman 2018).

Growth of personal genomic data has been limited by bottlenecks in computational requirements and server capacity (O’Driscoll et al. 2013). The NIH and several other institutions are moving toward cloud-computing-based services in order to overcome these bottlenecks (Patterson 2018; Navale and Bourne 2018; GovernmentCIO 2019). However, cloud-based storage and data analysis tools present security concerns, as they are based on a centralized architecture and are therefore vulnerable to single-point-of-failure losses (Ozercan et al. 2018). They also require trust in a third-party company (i.e. Google or Amazon), which may not be desired. These are critical problems; as genomic data becomes increasingly integral to our understanding of human health and disease, its integrity and security must be a priority when providing solutions to storage and analysis. Corruption, change, or loss of personal genomes could create problems in patient care and research integrity in the future. Another issue associated with data storage is data ownership. When an individual purchases a sequencing service from a company such as 23andMe, they are often giving that company the right to monetize their genomic data by selling it to pharmaceutical companies (Rosenbaum 2018). Ideally, individuals would retain ownership and be able to benefit from their own data (Grishin et al. 2018). Yet, the storage infrastructure to achieve this goal is lacking.

An ideal implementation of personal genomic data storage would 1) protect from loss and manipulation, 2) provide appropriate access to clinicians and biomedical researchers, and 3) allow the individual control over their own genomic data. In recent years, Blockchain has emerged as a potential solution that would achieve these requirements. Blockchain has several key properties which could make it an ideal solution, including a decentralized, distributed architecture, and cryptographic protocols that yield immutability. Already today, there are multiple personalized-medicine startups that aim to use blockchain to improve genomic data storage.

Proposed in 2008 as a global cryptocurrency network (Nakamoto 2008), blockchain is now used to solve a variety of problems. Blockchain consists of an append-only data structure shared in a decentralized, distributed network, which can be public or private. The network relies on a consensus mechanism to add to the chain. Most widely known is proof-of-work (PoW), in which nodes in the network participate in a mining competition. Others are proof-of-stake (PoS) and proof-of-authority (PoA). Public blockchain networks typically make use of PoW, which is suitable for large networks of individuals of unknown identity and intent. On the other hand, private blockchain networks consisting of individuals with known, semi-trusted identity tend to use PoA. A previous study highlights the specific benefits of blockchain for biomedical research applications (Kuo et al. 2017). These include decentralized management, immutable audit trail creation, data provenance, robustness and availability, security and privacy. A recent review points out possible use cases of blockchain in genomics research. These include distributed computation, data storage and distribution, voting on standardization protocols, crediting data ownership, and infrastructure for large scale organizations such as GA4GH, ELIXIR, TCGA, and ICGC, which must implement rules and regulations (Ozercan et al. 2018).

There are a few different blockchain platforms one might consider for developing a data-storage application. One is Bitcoin, a public cryptocurrency network. However, here we are concerned with private blockchain networks, as genomic data is sensitive and should only be shared with a set of individuals or institutions (e.g. sequencing centers, physicians, biomedical researchers) (Figure 1a). Whereas anyone in the world can participate in the public Bitcoin blockchain, only permissioned individuals can sync a private blockchain. Furthermore, Bitcoin supports only simple transactions and transfers of small amounts of data (~80 bytes) from one user to another. Another is Ethereum, which supports more complex transactions via Smart Contracts, self-executable on-chain programs which write to their own storage. Ethereum is a public network, but permits the creation of private networks. One of the most prominent platforms for private-blockchain development is MultiChain, a Bitcoin-like platform. Different kinds of blockchains are suitable for different use cases. For example, the Bitcoin blockchain is perfectly suitable for cryptocurrency exchanges. It simply acts as a ledger of transactions between different accounts. However, for more complex transactions or storage protocols, Ethereum becomes more suitable. MultiChain does not permit on-chain computation as Ethereum does, but it has several features that make it especially suitable for data storage use cases. Both the MultiChain platform and Ethereum Smart Contracts have been previously used in medical genomics applications, but in limited settings (Gursoy et al. 2020; Pattengale and Hudson 2020; Ozdayi et al. 2020; Ma et al. 2020; Gursoy and Brannon et al. 2020).

**Figure1.**
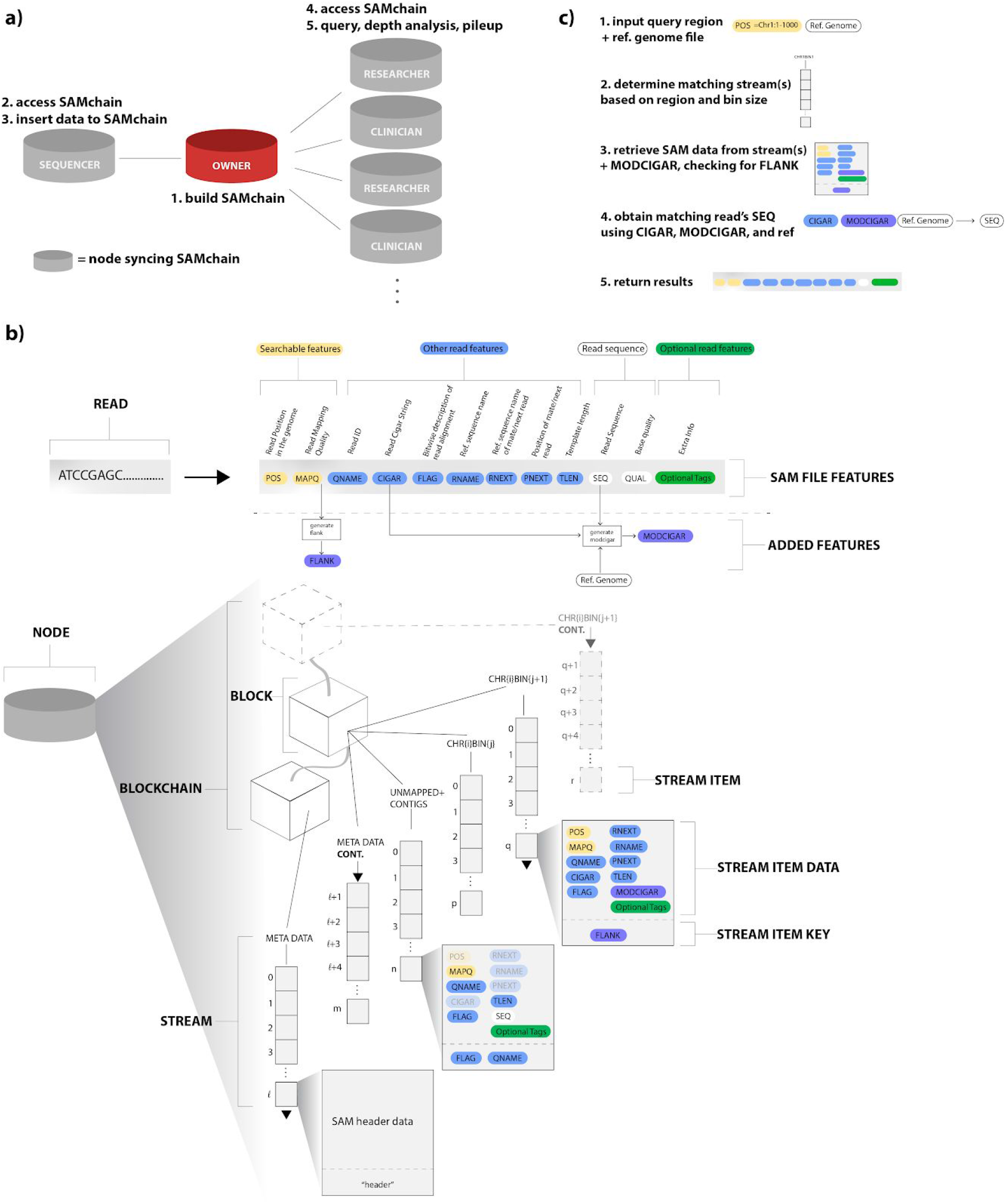
SAMchain design and implementation. (a) Overview of the SAMchain network ecosystem. The network consists of owner, sequencer, clinician, and researcher nodes. The owner node builds the SAMchain, the sequencer node accesses it and inserts SAM data to the chain. Clinician and researcher nodes access the SAMchain and analyze the on-chain SAM data. (b) Details of data storage in SAMchain. A read is typically stored in a SAM file containing several features. Our data structure is organized by genomic location. A single stream, named metaData, contains all of the header data and other chain info. Many other streams serve as bins by genomic location, and hold the SAM feature data and MODCIGAR. A FLANK feature is used to indicate whether a read’s position spans two consecutive bins. Stream items correspond to a single read. A single stream, named unmappedANDcontigs stores unmapped reads and contigs. (c) Overview of the query process. Upon querying a genomic location, our algorithm searches through the binned streams to obtain the SAM data and MODCIGAR features corresponding to the specified location. This data, in combination with a reference genome, yields a complete SAM read. Our algorithms and stream-based data structures are built on top of MultiChain, which provides the underlying blockchain, stream design, and network configuration.

Previous reviews outline the current status of commercial and academic proposals that use blockchain technology to improve genomic data sharing (Ozercan et al. 2018; DeFrancesco and Klevecz et al. 2019). Among these platforms are CrypDist, Zenome, Nebula Genomics, the Cancer Gene Trust, and Encrypgen/Gene-Chain. Each of these platforms utilizes blockchain for different aspects of the genomics data storage and sharing process. For example, Zenome makes use of Ethereum Smart Contracts to facilitate access to genomic data files and exchange of ‘ZNA tokens’, cryptocurrency which allow individuals to be compensated for their genomic data (Ozercan et al. 2018; Kulemin et al. 2017; Zenome.io 2017). Nebula Genomics also uses Smart Contracts to communicate between nodes in the network, survey participants, and facilitate data access permissions (Grishin et al. 2018). However, due to the difficulty of storing large data on blockchains, many of these companies store the genomic data elsewhere, such as in Blockstack or InterPlanetary File System (Grishin et al. 2018; Currie 2018). For example, CrypDist uses a custom Blockchain to store *links* to genomic data files (such as reference-aligned BAM files), which are actually stored in AWS data buckets (Sahin 2017). Because these platforms do not actually store genomic data ‘on-chain’, they are missing a key benefit of blockchain: high-integrity, secure data storage. Storing links to data in a blockchain can be useful in some cases, for example, if it is important to keep an access log for a particular dataset (Gursoy et al. 2020). Yet, it does not secure the data itself, nor does it prevent it from being altered, as it is stored somewhere else entirely. On the other hand, storing genomic data on-chain maintains the integrity and security of the data (Kuo et al. 2017). Additionally, while many existing platforms provide network architectures for storing genomic data, few offer solutions for performing computation on the data stored in the network. This is a critical gap in the technology; not only do clinicians and researchers need access to high-integrity, raw genomic data, but they also require secure tools for querying and streaming the data. This is likely due to a central caveat of blockchain technology: the inefficiency of storing and querying data due to the potential for chains to reach large sizes. The storage space and computational power required by blockchain is greater than a centralized database application due to the redundancy of storage and network verification protocols. The decentralized system also creates a higher latency (delay in data communication) during storage and retrieval of data. Additionally, transactions in the blockchain network require a cryptographic consensus verification, which makes them slow to publish data to the chain (Nakamoto 2008).

In this study, we present an open-source, proof-of-concept private blockchain network, which allows efficient storage and retrieval of raw genomic reads, often stored in a reference aligned format (sequence alignment map (SAM) files) (Li et al. 2009; GA4GH). To overcome the challenges described above, we developed novel data structures based on nested database indexing, file-format modifications and compression techniques with the open-source blockchain API MultiChain (Greenspan 2015). We made use of their ‘data stream’ feature, which allows users to create multiple key-value, time-series, or identity databases that can be used for data-sharing, time-stamping, and encrypted archiving (Greenspan 2015) (see Supplemental Information). We provide two tools: the first is SAMChain, which allows users to create a chain and insert SAM data into it, and the second is SCtools, which provides functions such as (a) querying, (c) depth analysis, (d) pile-ups for variant calling, and (e) re-creating SAM files and their derivatives (such as BAM and CRAM files).

## Results

### MultiChain provides the most advantages to the SAMchain use case

We first evaluated the advantages of major blockchain platforms, Bitcoin, Ethereum, and MultiChain, for our particular use case (Table 1):

- Bitcoin is primarily designed to be a public cryptocurrency blockchain network, and does not offer an efficient way of indexing and querying stored data. Thus, we quickly ruled it out for our particular use case.
- Ethereum makes use of Smart Contracts, which are Turing-complete, on-chain programs that can perform many different functions (other than just sending value from one account to another). While Ethereum can be used as a distributed database, it is more suitable for scientific computation.
- MultiChain is perhaps more suitable for database development due to built-in features called “streams”. Streams are append-only, on-chain lists of data with key:value retrieval capability, making store and query functionality extremely easy. From a developer’s perspective, MultiChain also has advantages of centralized documentation and ease of implementation.

**Table 1.** Brief summary of the big-picture differences between the Bitcoin, MultiChain, and Ethereum and their suitability for SAMchain.

| Bitcoin | MultiChain | Ethereum |
| --- | --- | --- |
| - transactions only<br>- public +private networks (better suited for public)<br>+ on-chain data storage<br>- off-chain computation<br>- distributed documentation<br>- less commonly used as database | + streams-easy key:value retrieval<br>+ public +private networks (better suited for private)<br>+ on-chain data storage<br>- off-chain computation<br>+ centralized documentation<br>+ commonly used as database | + smart contracts-customizable programs<br>+ public + private networks<br>+ on-chain data storage<br>+ on-chain computation<br>- distributed documentation<br>- less commonly used as database |

Given that we wanted to develop a robust software usable by individuals, physicians, and researchers for storage, query, and computation, we developed SAMchain with MultiChain. While MultiChain does not currently allow for on-chain computation like Ethereum does, its natural capacity to be used as a database outweighed this disadvantage. We provide an Ethereum Smart Contract in our code base as an example of how one might store raw genomic data in a Smart Contract, which would permit on-chain computations on the genomic data. As Ethereum becomes more widely utilized for database use cases, SAMchain could be expanded to Ethereum. In Table 1, we summarize the big-picture differences between Ethereum and MultiChain.

### Private Blockchain Network

We envisioned a network of sequencers, owners, clinicians, and researchers, each syncing SAMchain (Figure 1a). The owner node initializes a SAMchain, including the data streams which will store the SAM data. The sequencer node generates the SAM data and requests access to the data owner’s SAMchain. The owner grants access, allowing the sequencer to push the owner’s data to their chain. The clinician and researcher nodes, upon making contact with the owner through other means, may also request access to the owner’s SAMchain and make use of the SCtools modules to analyze the data. In this scheme, the owner may change the permissions of SAMchain at any time.

### SAMChain’s design provides advantages over traditional blockchain data storage methods

Next we considered how best to configure data storage in SAMchain. The naive way to store data in a bitcoin-like blockchain would be to append small amounts of data to each transaction using OP_RETURN, a script opcode allowing the sender to send a small amount of data (which varies between platforms) with their transaction. The OP_RETURN data is mined into a block along with the rest of the transaction. The data that is pushed using OP_RETURN is indexed in a transaction with a unique identifier, often called “ref” in a transaction. Each transaction can hold data around ~80 bytes, which means genomic data needs to be stored within numerous transactions. The unique identifiers need to be stored somewhere separately, as one can retrieve data only with ref. As one can imagine, querying the genome with a specific position, or doing a pile up analysis with OP_RETURN is extremely difficult and inefficient due to lack of needed data structures (Sward et al. 2018). In addition to storing the data embedded in transactions, Multichain offers another data structure to store the data on the chain, called “data streams.” Each stream item is represented by a blockchain transaction, which is mined and validated by every node like any other transaction (Greenspan 2016). The embedded key:value property of the streams allow efficient retrieval of the data on the chain, because when a node in a MultiChain network subscribes to a stream, it indexes the stream’s content in real time in order to enable efficient retrieval by keys (MultiChain 2020). The naive way to store data in a MultiChain stream would be to push all the data to a single stream, and query from it. However, as shown in Figure 3 Panel b, for our use case retrieval time increases when a stream contains too many items, as the query algorithm must check through each item to determine if it matches the queried position or range of positions. Were each genomic read defined by a single point position, we would not have this problem; we would simply store point positions as keys. However, because a read is defined by a range of positions, and there is overlap between reads, our query algorithm must actually check that a given read overlaps with the range or point queried, which is not possible in a traditional key:value stored dictionary (i.e streams in MultiChain). To achieve this kind of query with efficiency, we created streams binned by genomic location. This increases the time it takes to create a SAMchain (which is only done once), as many streams must be created, but significantly decreases the query time, making SAMchain a viable way of storing raw genomic reads on a blockchain. In Figure 3 Panel d, we show the advantage in query time gained by the SAMchain design compared to the single-stream method, which is up to 25-fold. In Table 2 we summarize the contributions of MultiChain to our paper, vs. those of SAMchain itself.

**Table 2.** Contributions of MultiChain vs. SAMchain. The novel method presented in this paper is SAMchain, a tool that is built upon the functionality of the MultiChain blockchain platform/api. Here we summarize how the contributions of SAMchain differ from those of MultiChain.

| **MultiChain** | vs. | **SAMchain** |
| --- | --- | --- |
| provides underlying blockchain protocol |  | utilizes MultiChain blockchain as a tool to store SAM data |
| provides built-in stream feature |  | indexes streams by genomic location |
| provides commands to retrieve stream data |  | combines stream items with reference genome to return complete SAM data |

### SAMChain provides a platform for storing next-generation sequencing data in blockchain

It is notoriously difficult to store large amounts of data in a blockchain, due to network latency and storage redundancy (Kuo et al. 2017). Thus, to store raw genomic data on-chain, we first needed to manipulate the information content for efficient storage. Whereas a SAM file stores a read’s sequence and the quality string of the read, SAMchain stores only the difference between a read and a reference genome. This design was inspired by the CRAM file format, in which a genomic reference file is optionally used to describe the difference between the aligned sequence and the reference sequence (Hsi-Yang Fritz et al. 2011). However, CRAM is a columnar file format composed of containers while SAMchain stores data in plain text. Our manipulation consists of storing a new data field that we refer to as the ‘modcigar’, a string containing the sequence data that differs from the reference (e.g. insertions). We show that there is a ~2 fold reduction in the storage with this manipulation (Figure 2a). Next, we designed the SAMchain data structure using MultiChain streams. As defined by MultiChain, streams are ordered lists of items, each with a publisher (who digitally signed the item), a set of keys (to be used for retrieval), some data (which is embedded on-chain in a transaction), and some meta data (about the transaction and block corresponding to the item). As we discuss in the section above and show in Figure 1X, querying data by genomic position from a single stream would be very time-inefficient. To address this problem, our code creates several streams binned by genomic position based on an input bin size. For example, the length of human Chromosome 1 (build GRCh38) is 248,956,422 base pairs. If the user were to set the bin size to one million base pairs, then 248,956,422/1,000,000 = 249 streams would be created for Chromosome 1, named chr1stream{j}, where j ranges from 1 to 249. In these streams, we stored the SAM features in the data field, and a feature called “flank” in the key field, which indicates whether a read’s coordinates span one stream (flank=0) or two (flank=1). The flank feature is necessary because the position of some reads will naturally span two consecutive streams. This design allows our query algorithm to know exactly which streams to search during a query, and to search through streams with fewer items. We confirmed that retrieval time in MultiChain is based only on the number of entries in a given stream (and unaffected by other streams on-chain) (Figure 3a). Our code also creates a metaData stream to store the file header, and an unmappedANDcontigs stream to store any unmapped reads and contigs in the case that they can be realigned in the future. The design of the SAMchain data structure is shown in Figure1b.

**Figure 2.**
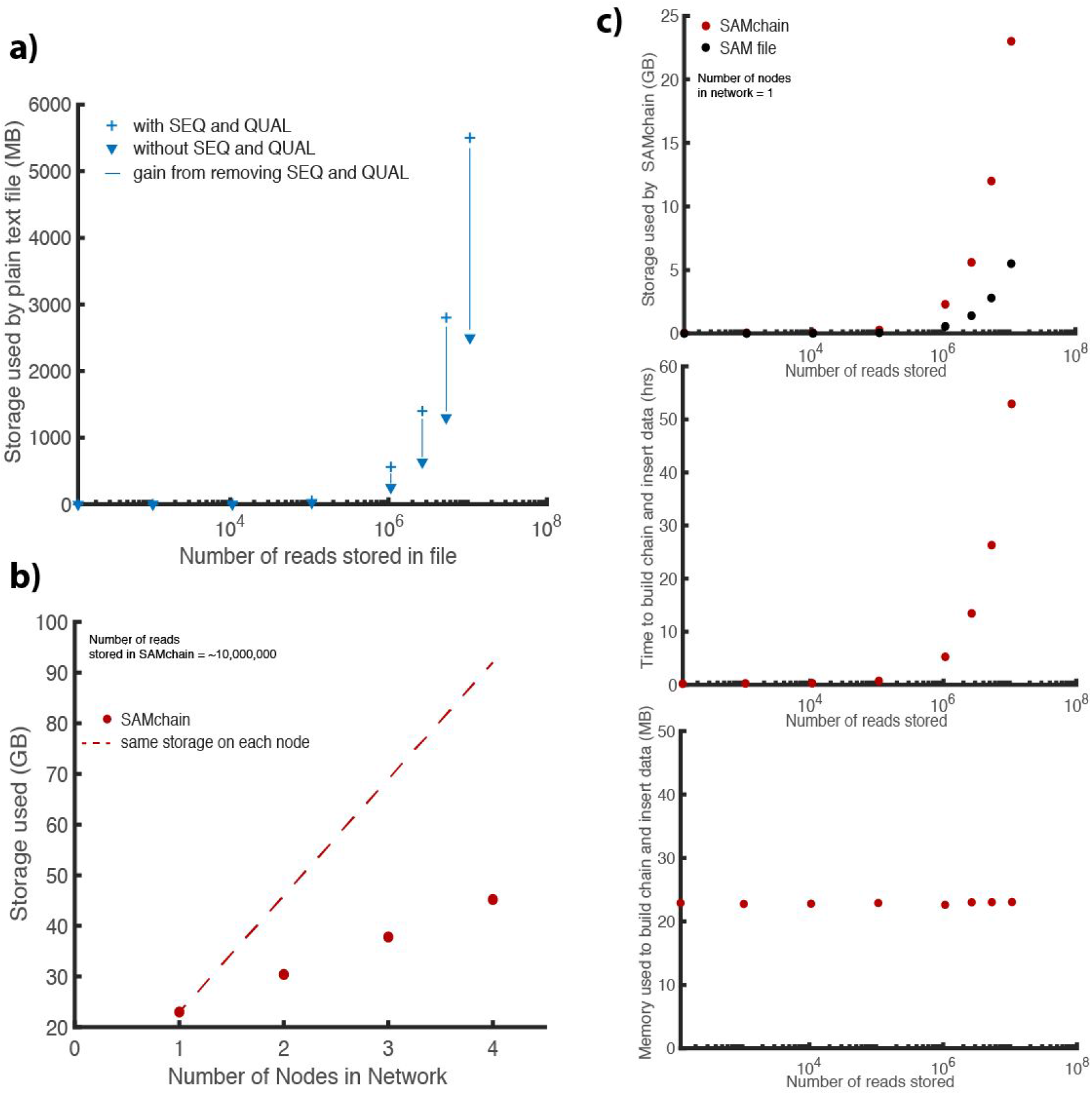
SAMchain performance. **a)** Before inserting SAM data to a SAMchain, we remove the sequence and quality strings. In plain text, we measured the storage gain by removing these fields. **b)** The total network storage required for a SAMchain storing ~10 million reads **c)** Storage (compared to a SAM file), time, and memory used to build a SAMchain.

**Figure 3.**
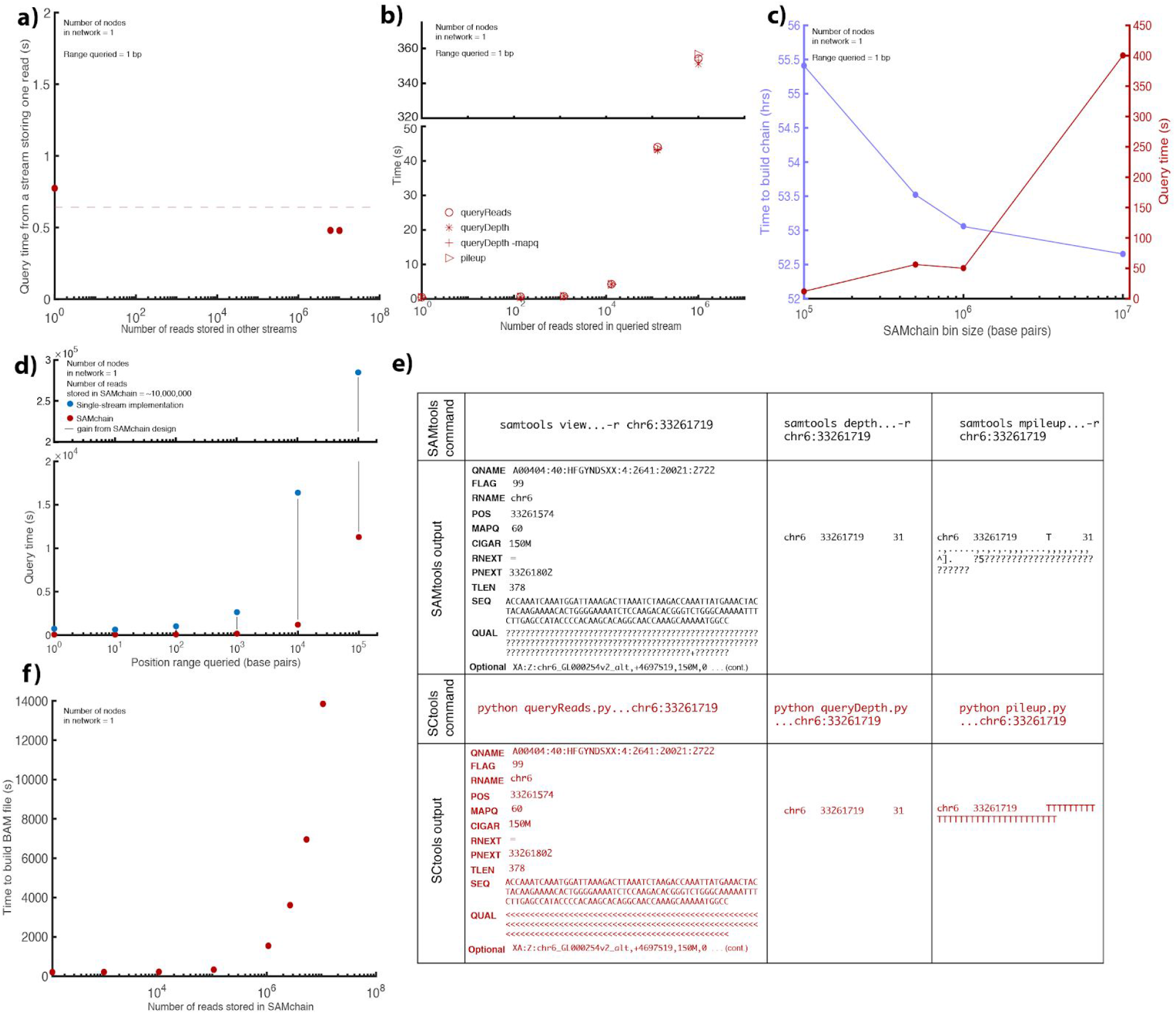
SCtools performance. (a) Query time from a MultiChain stream depends only on the number of entries in that stream, i.e. is not affected by the number of reads stored in other streams in the chain. (b) In a single-node SAMchain network, we measured the time performance of queryReads, queryDepth, queryDepth AND mapq, and pileup for one-bp queries. Each module performs comparably, increasing linearly as the number of reads stored in the chain increases. (c) We measured the performance of queryReads for a SAMchain storing ~10,000,000 reads at different bin sizes. SAMchains with smaller bin sizes yield faster query time, but take longer to build. (d) We measured the effect of the range queried on performance time. Larger ranges are shown to yield longer query times. We showed the times of SAMchain (red) compared to a naive, single-stream implementation (blue). (e) We checked the output of each SCtools module compared to the equivalent SAMtools module.

We measured time, memory, and storage requirements of building a SAMchain, and the time requirements of SCtools. All the tests described in Results are done using an alignment file curated from the high-coverage whole genome sequencing data of individual NA12878 from the 1000 Genomes Project as input data. For speedy testing, we constructed a SAM file with one disease-causing locus of roughy 1-5 million base pairs per chromosome. The details of these loci are given in SI Table 2. While we ran performance evaluation with DNA sequencing data, SAMchain and SCtools are compatible with any next generation sequencing data, including functional genomics data. For example, SCtools can be utilized to query variants in a gene or the depth distribution of exons of interest on raw RNA-Seq reads.

For a fixed bin size (1 million base pairs) and varying number of input reads, we evaluated the time, storage, and memory requirements of building and inserting data to a SAMchain. As shown in Figure 2c, the storage required per node by SAMchain is approximately 5-fold greater than that required by a SAM file. For example, our input SAM file containing ~10 million reads requires 5.5 GB. Building the corresponding SAMchain in a one-node network requires ~25 GB. As shown in Figure 2b, storage requirements of a SAMchain increase with increasing nodes in the network, as each node stores redundant data. For a chain storing ~10 million reads, each additional node required 7.4 GB. The node that inserts the SAM data to the chain will require the most storage because MultiChain keeps a “wallet” directory, which stores transaction data especially relevant to the local node~(see SI section 2.1 for details). In Figure 2c, we measured the time and memory it takes to build and insert data to a SAMchain as a function of the number of reads in the input SAM file, which show linear and constant trends, respectively. To build and insert 10,00,000 reads to a SAMchain, it takes ~55 hours. However, this must only be done once.

### SCtools can query NGS properties directly from SAMChain

We developed SCtools to query reads and other NGS properties from a SAMchain. Specifically, we developed four modules: queryReads, queryDepth, pileup, and buildBAM. queryReads queries a SAMchain based on one of four SAM features and outputs reads in SAM format. Currently, the feature we use for querying is genomic location. queryDepth performs depth analysis on an input range of genomic coordinates. pileup performs pileup analysis on an input range of genomic coordinates. buildBAM reconstructs a BAM file from a SAMchain. To show the scalability and performance of these modules, we first showed that the time to perform a point query depends only on the number of reads in the queried stream and is not affected by reads stored in other on-chain streams (Figure 3a). Next, we measured the time requirements of each SCtools module as a function of the number of reads stored in the queried stream (Figure 3b). The modules show comparable time efficiency, increasing linearly with the number of reads stored in the queried stream (Figure 3b). We also checked the effect of “AND” queries, that is, filtering a depth query by mapq score (Figure 3b, ‘+’ icon). We found that this type of query performed just as well as the other modules. In Figure 3 Panel c, we investigated the impact of changing the SAMchain bin size for a fixed number of reads stored (~10 million). We found that increasing the bin size increases the query time, which was expected because a larger bin size also contains a higher number of reads relative to a smaller bin size. However, increasing the bin size also decreases the total number of streams, which reduces the time it takes to build the SAMchain (Figure 3c). In Figure 3 Panel d, we measured query time for a SAMchain storing a fixed number of reads (~10 million) as a function of the range of genomic coordinates queried. We found that query time increases linearly with increasing range queried. In Figure 3 panel e, we show the output of each SCtools module compared to that of the comparable SAMtools function. And in Figure 3 panel f, we measured the time it takes to build a BAM file from the read data stored in a SAMchain, and found that the time increases linearly with increasing number of reads stored in the chain.

### VCFChain and VCFquery modules provide faster options for direct query of genomic variants

There is significant value in storing and sharing aligned genomic reads, as opposed to just the genomic variants called from the aligned reads. Access to raw reads is important because if a new reference genome build is available, the reads can be re-aligned and new variants may be observed, thereby maximizing the utility of the data. Thus, our primary goal was to create a blockchain storage system for SAM files. However, there can also be value in sharing variant data (VCF files). Thus, we also developed VCFchain and a VCF query module. VCFchain stores data from a VCF file on chain, and VCF query can query variants from the chain by genomic position, along with rsid and/or genotype. To test these modules, we used a VCF file from a consented individual in the ENCODE data portal. We chose this vcf because it does not only contain SNPs and small indels but also a full set of structural variants. In Figure 4, we show the performance of building a VCFchain and VCFquery. As shown in Figure 4a, the storage required for a one-node network per node by a VCFchain storing ~6.5 million variants is approximately 40 GB. As more nodes are added to the network, total storage increases. In this case, each additional node requires 14 GB. In Figure 4b, we measured the storage, time, and memory requirements of VCFquery in a one-node network with an increasing number of stored variants. In Figure 4c, we measured time requirements of VCFquery for position queries, along with position + rsid and position + rsid + genotype queries. Figure 4d shows the output of VCFquery for a given read compared to that of bcftools view.

**Figure 4.**
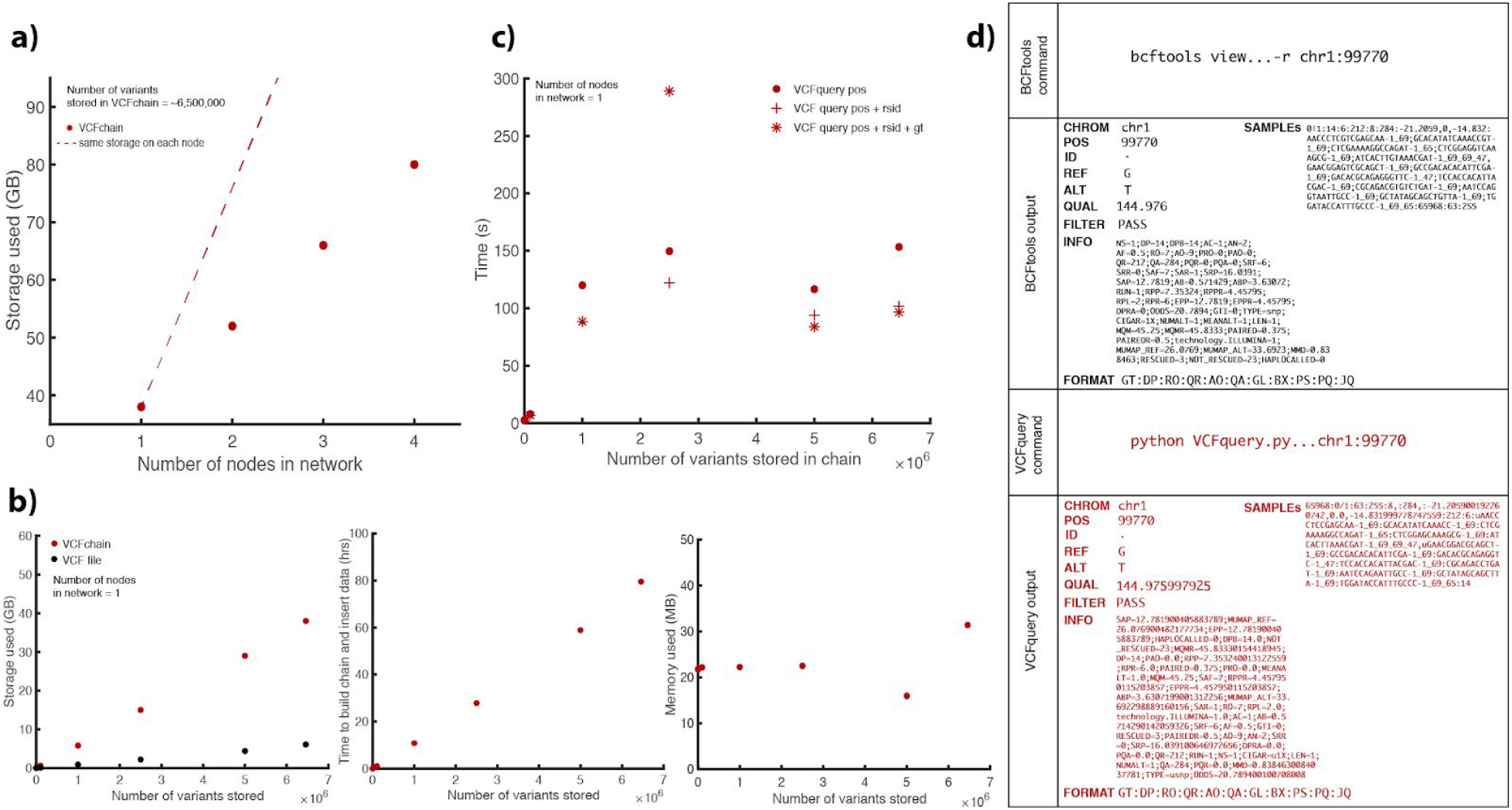
VCFchain and query performance. (a) The total network storage used by a VCFchain (storing 6458146 variants) as a function of the number of nodes in the network. (b) The storage, time, and memory requirements of building and inserting data to a VCFchain in a single-node network as a function of the number of reads stored in the chain. Storage is compared to that used by a VCF file. (c)The time requirements for VCFquery as a function of the number of reads stored in the chain, compared to retrieval time from a VCF file using bcftools. (d) Output of VCFquery compared to that of bcftools for a given read.

Our VCFchain infrastructure could also be used to store and share variants from a cohort of individuals. For example, one could build upon our code base to create a VCFchain with the somatic variants of TCGA dataset stratified by cancer type.

### Comparison to existing blockchain applications for genomics data

In addition to comparing SAMchain and SCtools performance to that of SAMtools, we also looked into the details of other genomic data storage platforms using blockchain. Because the role of blockchain in these platforms is qualitatively different from that in SAMchain, we are not able to make quantitative comparisons. For example, storing links in a blockchain to data stored elsewhere is fundamentally different than storing the data itself on-chain. Instead, we highlight the major difference between the platform and SAMchain in Table 3. Note that the information about each platform is derived from the limited knowledge provided by the company websites and whitepapers and is presented here to the best of our knowledge. We identified four companies and/or projects which use blockchain in the context of genomic data storage: CrypDist, Zenome, Nebula Genomics, and the Cancer Gene Trust. Another is Encrypgen/Gene-Chain, but their limited documentation prevents any comparison with SAMchain. While each of these four platforms uses blockchain for some aspect of their network ecosystem, none uses it to store raw genomic reads on-chain. CrypDist stores links to data, which appear to be stored in AWS buckets (Sahin et al. 2017); Zenome uses Ethereum Smart Contracts to facilitate transactions of genomic data, but appears to store the data off-chain in a distributed file storage system (Zenome.io 2017; Ozercan et al. 2018); Nebula Genomics uses Ethereum Smart Contracts to facilitate communication between nodes, and Blockstack to facilitate data storage, but Blockstack stores the data off-chain, either on a local drive or in the cloud (Digital Ocean, S3, Dropbox) (Grishin et al. 2018; Blockstack docs; Defrancesco and Klevecz 2019); and finally, the Cancer Gene Trust stores data off-chain in InterPlanetary File System and raw genomics data locally, and uses Ethereum Smart Contracts to store references to the data files (Cancer Gene Trust 2018; Currie 2018; Defrancesco and Klevecz 2019; Ozercan et al. 2018). To the best of our knowledge, SAMchain is the first proof-of-concept project to store raw genomic reads on a blockchain, on-chain. By embedding the data in the chain, SAMchain aims to preserve the integrity of the data by taking advantage of blockchain protocols designed to preserve the integrity of cryptocurrency transactions.

**Table 3.**
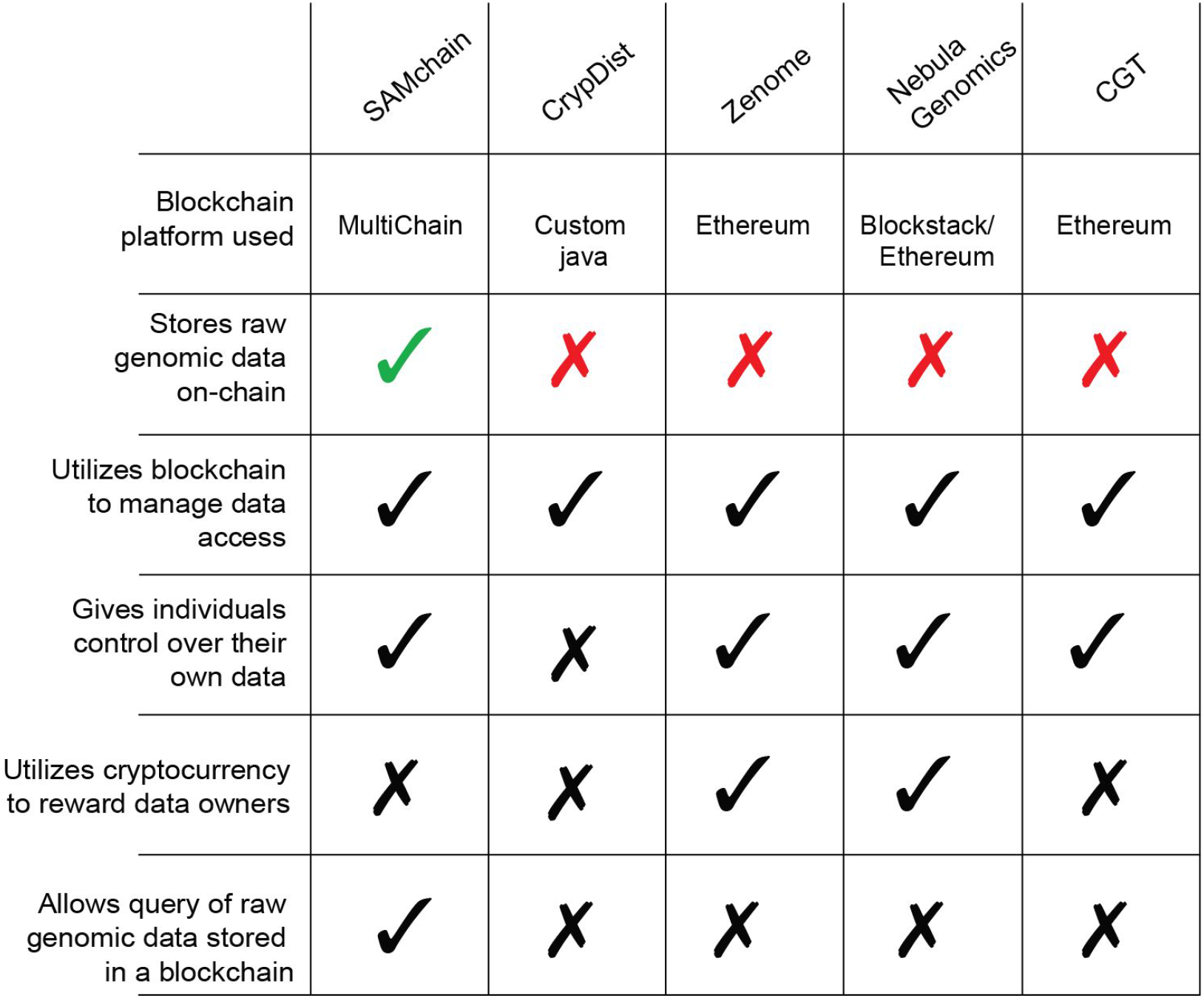
Side-by-side comparison of existing blockchain genomics data storage platforms and SAMchain. Importantly, SAMchain is the only one that stores raw genomic data on-chain, leading to high-integrity data storage.

|  | SAMchain | CrypDist | Zenome | Nebula<br>Genomics | CGT |
| --- | --- | --- | --- | --- | --- |
| Blockchain platform used | MultiChain | Custom java | Ethereum | Blockstack/<br>Ethereum | Ethereum |
| Stores raw genomic data on-chain | ✓ | ✗ | ✗ | ✗ | ✗ |
| Utilizes blockchain to manage data access | ✓ | ✓ | ✓ | ✓ | ✓ |
| Gives individuals control over their own data | ✓ | ✗ | ✓ | ✓ | ✓ |
| Utilizes cryptocurrency to reward data owners | ✗ | ✗ | ✓ | ✓ | ✗ |
| Allows query of raw genomic data stored in a blockchain | ✓ | ✗ | ✗ | ✗ | ✗ |

## Discussion

We envision a real-world scenario in which individuals create private blockchains to store their personal genomes to share with their healthcare providers and biomedical researchers. Simply with ssh access, healthcare providers and associated genetics researchers can stream or query patients’ genomes. This reduces not only the risk of data corruption, but also non-permissioned access to private data. Blockchain provides immutability such that the data cannot be altered, whether intentionally or accidentally.

Our framework is the first open-source application to allow querying and streaming of genomic data from blockchain to the best of our knowledge. This is a substantial improvement over the current biomedical applications of blockchains. To address privacy concerns, our framework may be extended to store encrypted data in the data streams. One could even encrypt the data homomorphically, allowing direct computations on the encrypted data (MultiChain 2020). However, this would add storage and computation overhead to the solution.

While the main benefit of using blockchain for data storage is data security and integrity, blockchain also makes it easy to append data to large data files. For example, in the cases of SAM files, if a user wishes to add to the data, one could create data streams for these additions (since it is a private chain, only the owner and the permissioned users of the chain would have permission to make these changes). Thus, the data owner does not have to deal with opening large-data files, modifying them, and re-indexing them, which creates costly network traffic. Searches by genomic location could also check the new data streams, to determine if the owner has appended any changes to the data. Furthermore, the stream format lends itself well to storing reads; as discrete, independent items, reads naturally fit into stream items. To make computation completely on-chain, one could adapt SAMchain and SCtools for Ethereum. To demonstrate how this might be done, we provide a sample Smart Contract in our SAMchain code repository. As Ethereum becomes more and more suitable for database development, this will be an interesting future direction. Another future direction is to create dictionaries from the SAM files, in addition to reference-based compression techniques, compatible with blockchain querying mechanisms in order to further reduce chain storage requirements.

Our blockchain solution can be generalized to other large-scale data storage and querying problems beyond SAM files. Data including but not limited to electronic health records, vcf files from multiple individuals, and somatic mutation datasets from cancer patients can be stored in blockchain using our indexing schemes, allowing for rapid and partial retrieval of the data.

## Methods

### MultiChain specifics

We designed SAMchain as a layer on top of MultiChain. MultiChain data streams make it possible for a blockchain to be used as a general purpose database. The data published in every stream is stored by all nodes in the network. Each data stream on a MultiChain blockchain consists of a list of items. Each item in the stream contains the following information, as a JSON object (Greenspan 2015): A publisher (string), key:value pairs (between 1-256 ASCII character, excluding whitespace and single/double quotes) (string), data (hex string), a transaction ID (string), blocktime (integer) and Confirmations (integer). When data needs to be queried or streamed, it can be retrieved by searches using the key:value pairs. Publishing a stream item to a data stream constitutes a transaction. When a node subscribes to a stream, it indexes the stream items in different ways to enable fast retrieval, and the index entry points to the transaction ID. Because of the peer-to-peer network architecture, stream items can arrive at different nodes in the network in different orders. MultiChain permits the user to set a network ordering parameter to be Local or Global (default is Global, used for SAMchain). Global ordering means that once the chain has reached consensus, all nodes see the same order in their streams. Transactions submitted to the network are time stamped via Linux timestamp. When a transaction happens, it is held in the memory pool. After mining of the transaction is complete, the transaction is added to a block. Each block has a maximum transaction size, i.e after a block reaches its maximum size or the time to create a block reaches its limit, the block is sealed and appended to the chain. This means a data stream in MultiChain can span multiple blocks based on the time of the transaction (i.e time of the publishing the data to the blockchain). New blocks are created according to the “target-block-time”, a parameter set upon initializing a chain. We used the default value, which is 15 seconds. We also used the default consensus mechanism, which is a round-robin schedule of miners (Greenspan 2015). Users can also turn on proof-of-work mining if they desire, but the security of MultiChain does not depend on the proof-of-work scheme. As outlined in the MultiChain whitepaper (Greenspan 2015, pg. 3), problems can arise if proof-of-work mining (used in the public Bitcoin blockchain, for example) is applied to a private or “institutional” setting, such as the “51% attack,” in which over half of the permissioned participants collude to alter the chain (Greenspan 2015). To resolve this issue, MultiChain makes use of a “mining diversity” parameter, which controls the number of blocks which may be created by a given user within a set time window. Tuning this parameter changes the proportion of the network that would need to collude in order to undermine the network (detailed on pg. 7-8 of Greenspan 2015). Therefore, even though MultiChain is a private network, immutability is achieved.

We designed SAMchain, SCtools, VCFchain, and VCFquery using MultiChain version 2.0.3 and Python version 2.7.16. Together, the SAMchain and SCtools repositories contain six modules: buildChain, insertData, buildBAM, queryReads, queryDepth, and pileup. VCFchain and VCFquery contain three modules: buildChain, queryPosition, and queryGT-RSID.

### SAMchain and SCtools design overview

We took an approach to maximize the efficiency of storing and querying data. Our goal was to store minimal data while indexing it in a creative way to allow rapid retrieval, thereby reducing the time and memory cost of analysis and increasing the utility of the stored data (Figure 1a). To achieve this goal, we manipulated a) data structures in data streams, and b) data to be stored in SAM files. For (a), we first separated mapped and unmapped reads from a SAM file. First, we created a data stream called metaData to store the header data and general information (bin size, number of streams, etc.) about the chain. We then created N streams (called chr_i-bin_j). Each of these N streams represents a bin of genomic coordinates. Based on the location of a read mapped on the human reference genome, we log the read names as data in chr_i-bin_j stream. Some reads will span two bins. In that case, we store the read in the bin to which the beginning of the read maps. We then add a boolean key to the chr_i-bin_j stream that we call FLANK (SI Figure 1). FLANK=0 indicates that the entire read is in that bin. FLANK=1 indicates that the read coordinates span two consecutive streams. The FLANK value tells our retrieval algorithm to search for a particular read in two chr_i-bin_j streams. Our query algorithm can retrieve the data in the chr_i-bin_j stream based on the queried location. Our code base allows developers to bin the data according to a desired feature that might be queried by users such as read names, mapping qualities, or alignment scores; these are the features stored as keys in the binned streams. Our implementation uses binning by genomic location, as it is the most commonly queried property for depth analysis or variant calling. Unmapped reads are stored in a separate stream called unmappedANDcontigs, but not in the chr_i-bin_j streams. For (b), we were inspired by the data compression techniques in CRAM files (Hsi-Yang Fritz et al. 2011, Gursoy et al. 2019), and stored the difference between the read and the reference sequence in the chain instead of the sequences themselves to reduce the size of the data stored on-chain (SI Figure 6). With this approach, our implementation is able to regenerate the sequence of a read by using the reference genome and other features stored in the chain.

We developed SCtools to extract information from SAMchain for downstream analysis. We provide a code base that has the ability to query on a blockchain. The key:value property of the data streams in MultiChain2.0 together with the ability to query on multiple keys provides an opportunity to extract data from the blockchain without the need for costly calculations. Our query module can retrieve data from a chain based on the position in the reference genome(Figure 1c).

If a user queries the chain for reads mapped to a genomic region, our query module first finds the correct streams/bins containing that region. From the bins, it extracts the SAM data and MODCIGAR, and uses an input reference file to return the results. This approach reduces the query time significantly due to the following reason: Data streams do not allow range searches. If the data were kept in a single stream, then the query would have to iterate over the location range for every single stream item. With binned streams, the query is only done on the streams containing the relevant data.

Below we describe the functionality of each SAMchain, SCtools, VCFchain, and VCFtools module.

#### buildChain (owner node)

buildChain initializes a MultiChain blockchain and creates streams that will define the SAMchain. Three types of streams exist in SAMchain: 1) metaData, 2)unmappedANDcontigs, and 3) binned streams. metaData is a single stream which stores SAMchain settings (bin length, read length, and number of bins) and will eventually store the header from an input SAM file. unmappedANDcontigs is a single stream which stores the features from the input SAM file, except for the sequence and quality string, for unmapped reads and contigs. When parsing an input SAM file, the insertData module will use the FLAG feature to determine whether to put a read into the unmappedANDcontigs stream or a binned stream. Binned streams are a series of streams which map to a range of positions in the genome. buildChain divides each chromosome into N kb intervals (N is set by the developer) and creates a stream for each interval. We made this design choice to improve query efficiency. A user’s query leads to a specific binned stream (or set of streams), rather to all the data.

#### insertData (sequencer node)

insertData pushes data from an input SAM file to the relevant streams in an initialized SAMchain. First, insertData uses pysam to extract the header data from an input SAM file and pushes the header, line-by-line, to the metaData stream. Next, it uses pysam to extract the features of the reads in the SAM file, one at a time, and checks the read’s flag feature to determine whether it belongs in the unmappedANDcontigs stream, or a binned stream. It then pushes the read features to the appropriate stream as the data field of a single stream item. It checks whether the read’s position spans two streams. If it does, it stores that read in the stream mapping to its start position and stores “FLANK=1” as a key. If it does not, it stores “FLANK=0” as a key.

#### buildBAM (clinician/researcher node)

buildBAM rebuilds a BAM file from the data stored in a SAMchain. It first retrieves the header data from the metaData stream. Next, it retrieves the data from the binned streams and converts it to a tab-separated format. Using pysam, it extracts from an input reference file the sequence string and alters it based on the cigar. Finally, it uses pysam to write the read entry to an output BAM file.

#### queryReads (clinician/researcher node)

queryReads searches a SAMchain for reads that match an input region of interest in the genome. It first pulls information from the metaData stream about how the reads were binned during buildChain, and uses it to obtain the names of the stream(s) that correspond to the input genomic location. If the first stream in this list is not the first stream of a chromosome, it adds the stream name just upstream, in the case that a FLANK=1 read is present in that stream. Given these stream names, it uses built-in MultiChain commands liststreamitems and liststreamkeyitems to retrieve the items from those streams and check whether they match the region queried. Then, using the modcigar, it extracts the correct sequence from the reference genome and returns the results.

#### queryDepth (clinician/researcher node)

SAMtools provides a useful function to determine the sequencing depth for a queried location or all of the locations in the genome. queryDepth follows a similar algorithm to queryReads. However, after obtaining the read data, queryDepth must check the cigar values for each read in order to calculate depth taking into account information about insertions and deletions (for example, if a deletion occurs at a location queried in one of the reads, this should contribute +0 to the depth at that location). After calculating the depth values, queryDepth returns the results.

#### pileup (clinician/researcher node)

SAMtools provides a useful function to determine the pile-ups for a queried location or all of the locations in the genome. Pile-up files contain the number of reads that mapped to a location, the reference allele for that location, and the sequenced nucleotide in each read for that location. This allows users to visualize the genetic variation and calculate allele frequencies for the variants. pileup follows a similar algorithm to queryReads. However, after obtaining the read data, pileup must check the cigar values for each read in order to output pileup taking into account information about insertions and deletions. After doing so, pileup returns the results.

### VCFchain & VCFquery

#### buildChain

buildChain initializes a MultiChain blockchain and creates streams that will define the VCFchain. Two streams exist in VCFchain: 1) metaData and 2) allVariantData. metaData will eventually store the header from an input SAM file. allVariantData will store the features from the input VCF file, with genomic position, genotype, and rsid as the keys. All variants will be inserted to the allVariantData stream.

#### queryAND

Because positions in a VCF file are unique, queryAND retrieves VCF feature entries from the allVariantData stream using input genomic position as the key. It can filter data based on an input genotype and/or variant ID.

## Data Access

SAMChain and SCTools can be found at https://github.com/gersteinlab/SAMChain. The Ethereum smart contract, and VCFChain code can also be found in the github page. The SAM file (BAM) used in this manuscript can be found in http://homes.gersteinlab.org/people/gg487/samchain/. The vcf file used can be accessed at https://www.encodeproject.org/files/ENCFF907ASL/.

## Acknowledgements

We thank Dr. Mihali Felipe for the help with setting up the servers to run SAMChain in a multi node environment.

