## Supplemental Information for "Storing and analyzing a genome on a blockchain"

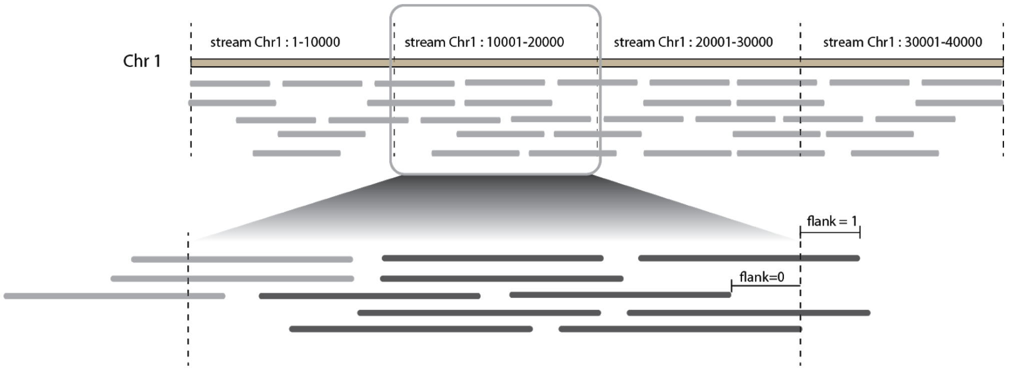


**SI Figure 1. Binning reads by position in streams.** To store read data from a sequence alignment file, we define several streams, each with a fixed position window. Some reads may span two of these streams. Thus, for each read we store a field called FLANK, which has value 0 if the data is in one stream, and value 1 if it spans two streams.

| | Key | Data |
|---|---|---|
| metaData | “chainInfo” | {<br>"json":{<br>"binLength":<binLength>,<br>"readLength":<readLength>,<br>"numBins":<numberBins><br>}<br>} |
| | “Header” | {<br>"json":{<br>"header":<headerText>,<br>}<br>} |
| chr{i}stream{j} | “FLANK={1 if hangs, 0 if not}” | {<br>"json":{<br>"QNAME":<readName>,<br>"FLAG":<flag>,<br>"RNAME":<chromosome>,<br>"POS":<position>,<br>"MAPQ":<mapq>,<br>"CIGAR":<cigar>,<br>"RNEXT":<next reference id>,<br>"PNEXT":<next reference start>,<br>"TLEN":<template length>,<br>"SEQ":<query sequence>,<br>"QUAL":<query qualities>,<br>"OP":<tuple of optional tags>,<br>}<br>} |

**SI Table 1**. Distribution of data across key and data fields of different types of streams.

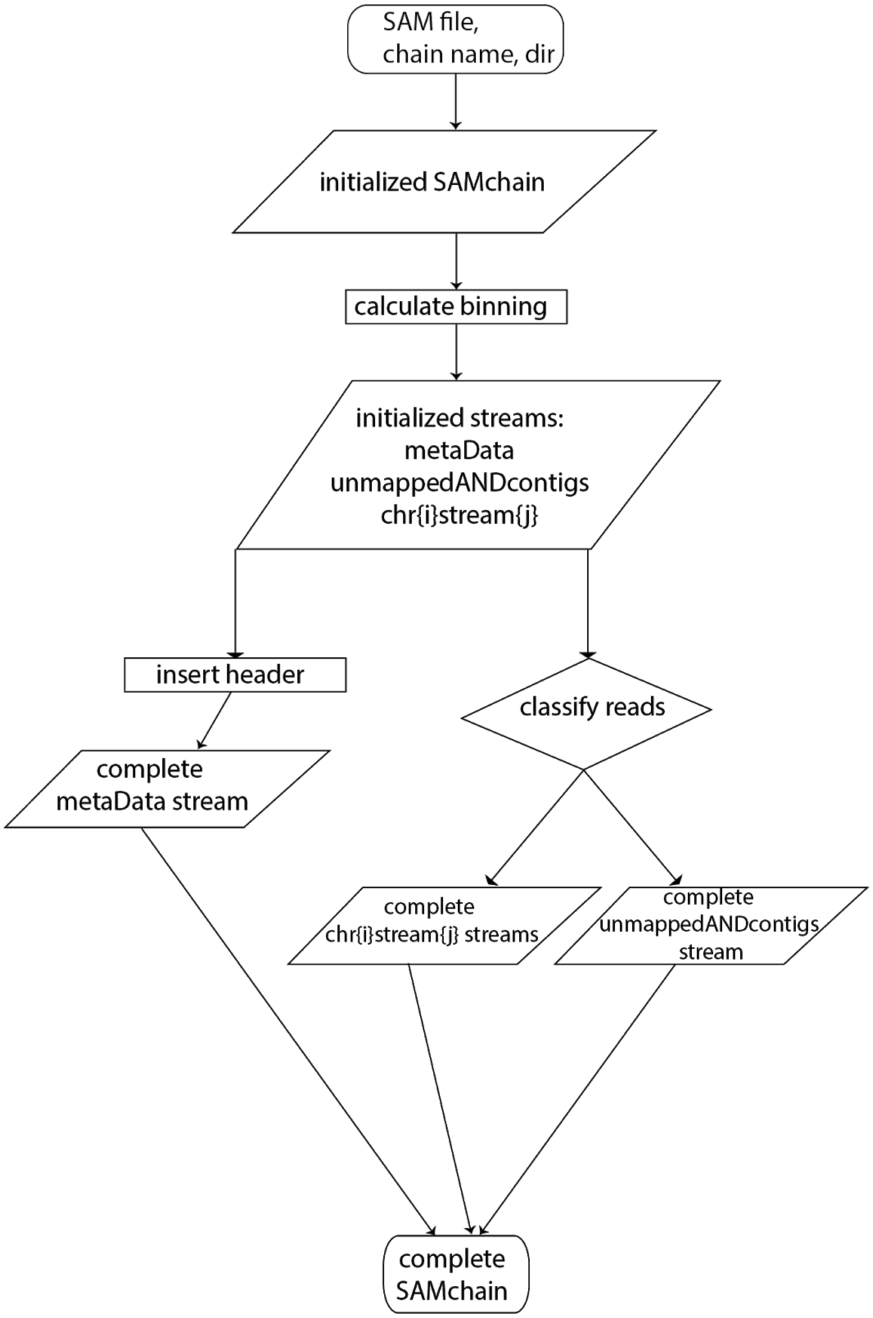


**SI Figure 2**. Flowchart for buildChain.py, a module which builds a SAMchain from an input BAM file.

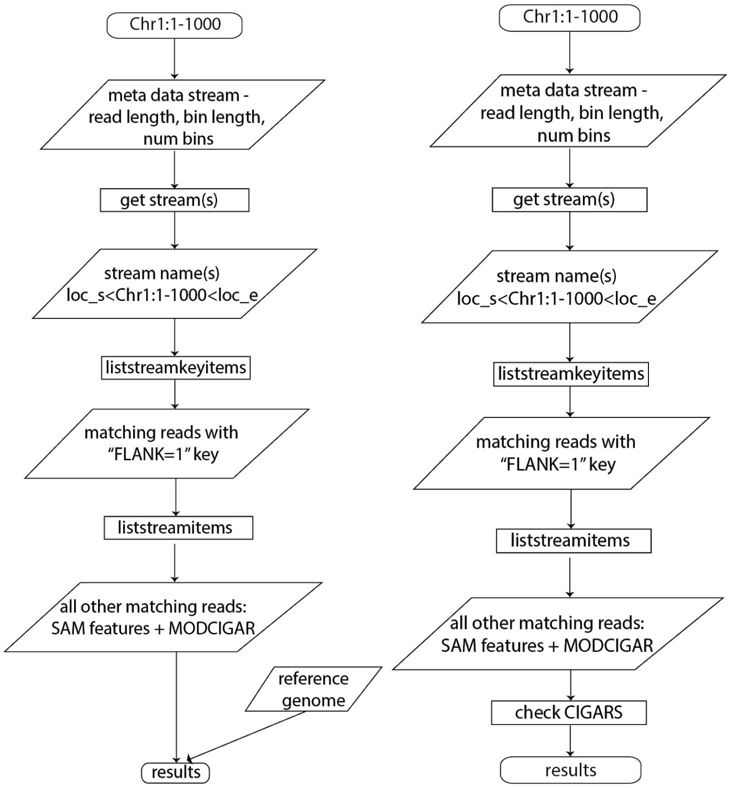


**SI Figure 3.** Flowcharts for queryReads.py (left), queryDepth.py (middle), and pileup.py (right), the core modules of SCtools.

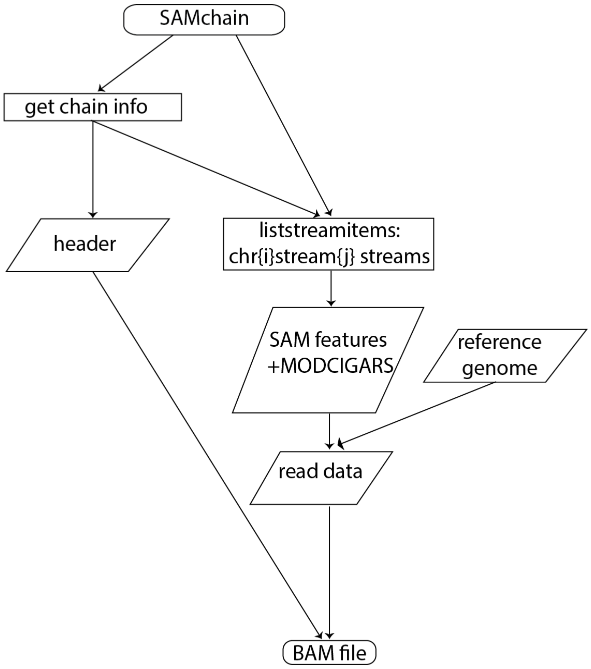


**SI Figure 4**. Flowchart for buildBAM.py, a module which rebuilds a BAM file from data stored in a SAMchain.

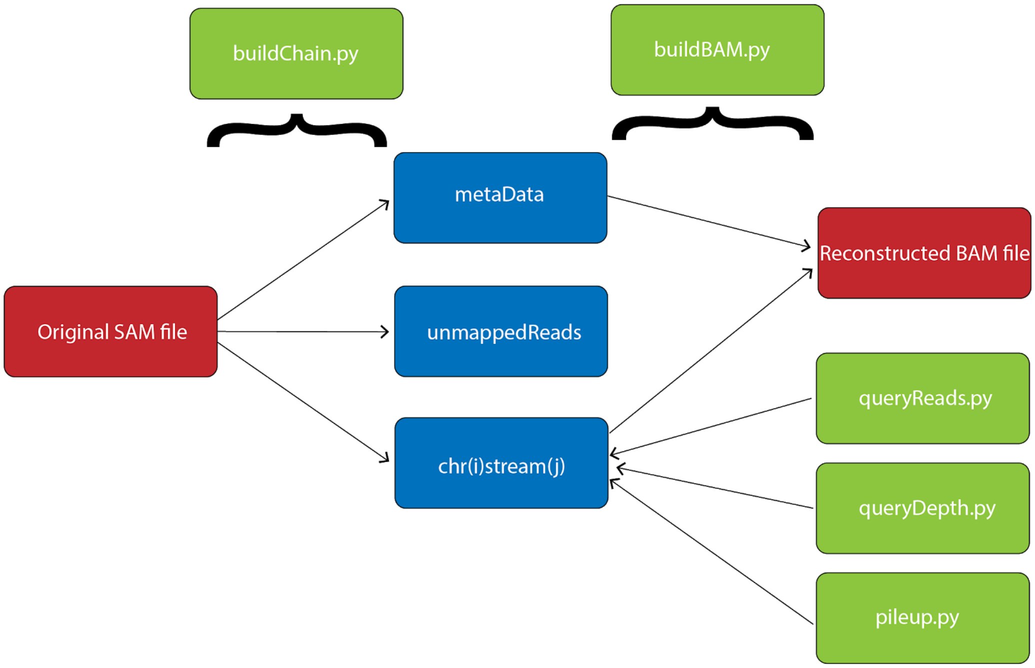


**SI Figure 5.** Overview of the SAMchain and SCtools organization and functions.

Genome used: HG00114, 1000 Genomes Project
Build: GRCh38
Source of loci information: https://www.ncbi.nlm.nih.gov

| Chromosome | Locus/Region | Coordinates | Length |
|---|---|---|---|
| chr1 | DAB1 | 56994778-58546726 | 1.5mil bp |
| chr2 | LRP1B | 140231423-142132463 | 1.9mil bp |
| chr3 | ROBO2 | 75906675-77649964 | 1.7mil bp |
| chr4 | 4p13 region | 41200001-44600000 | 3.3mil bp |
| chr5 | CDH18 | 19471296-20575886 | 1.1mil bp |
| chr6 | HLA locus | 28510120-33480577 | 4.9mil bp |
| chr7 | CNTNAP2 | 146116207-148420998 | 2.3mil bp |
| chr8 | CSMD1 | 2935353-4995035 | 2.0mil bp |
| chr9 | PTPRD | 8314246-10613002 | 2.2mil bp |
| chr10 | PCDH15 | 53802771-55629182 | 1.8mil bp |
| chr11 | DLG2 | 83455009-85628535 | 2.1mil bp |
| chr12 | ANKS1B | 98729904-99984773 | 1.2mil bp |
| chr13 | GPC5 | 91398621-92867237 | 1.4mil bp |
| chr14 | NRXN3 | 77979904-79868291 | 1.8mil bp |
| chr15 | AGBL1 | 86079620-87031476 | 0.95mil bp |
| chr16 | RBFOX1 | 5239752-7713343 | 2.4mil bp |
| chr17 | ASIC2 | 33013087-34156806 | 1.1mil bp |
| chr18 | DCC netrin 1 receptor | 52340172-53535899 | 1.1mil bp |
| chr19 | 19p13.13 region | 37790235-38403391 | 0.61mil bp |
| chr20 | MACROD2 | 13995516-16053197 | 2.0mil bp |
| chr21 | RUNX1 | 34787801-36004667 | 1.2mil bp |
| chr22 | DGS/VCFS region | 18924718-21111383 | 2.1mil bp |
| chrX | Dystrophin | 31119219-33339460 | 2.2mil bp |
| chrY | PAR1 | 10001-2781479 | 2.7mil bp |

**SI Table 2.** Details of the loci in the SAM file used for testing SAMchain.

### 1.1 Details of other blockchain platforms:

**CrypDist**: CrypDist is a program initially developed as an undergraduate senior project (Ozerkan et al. 2018). The code base for CrypDist is written in java and available on Github (Sahin et al. 2017). CrypDist does not make use of any widely used blockchain platform (such as Ethereum or MultiChain), but rather constructs a custom blockchain in java. The CrypDist blockchain stores links to data files, which appear to be stored in AWS buckets (Sahin et al. 2017). Importantly, the role of blockchain here is to store links and to log access to data rather than the data itself. This application is useful for research purposes when there are multiple NGS files from a cohort of individuals, as storing all the alignment files from the cohort in a blockchain is not feasible due to the storage requirements. This application aims to protect the integrity of metadata, while the security and integrity of the actual SAM files are not provided by blockchain. SAMchain differs from this application because 1) it stores data on-chain, 2) it gives individuals control over their data, 3) it is better suited for a personal genomics use case, that is, individual-level analysis rather than population-level analysis.

**Zenome**: Zenome is a company founded in 2017 (Kulemin et al. 2017; Zenome.io 2017). It appears that Zenome uses Ethereum Smart Contracts to facilitate ownership of and compensation for data (Zenome.io 2017). Nodes in the network can sell their data or services for "ZNA tokens," cryptocurrency convertible to ether (Ethereum currency). Raw genomic data is not stored in the Smart Contract. According to our understanding of their whitepaper and Smart Contract code, it appears that the data is stored off-chain in a distributed file storage system (Zenome.io 2017; Ozercan et al. 2018). The main differences between SAMchain and Zenome are 1) SAMchain stores NGS data on-chain to maximize data integrity, and 2) SAMchain does not utilize cryptocurrency.

**Nebula Genomics**: Nebula Genomics is a personal genomics company founded in 2018 (Grishin et al. 2018). According to their whitepaper, the Nebula system partitions genomes into overlapping, variable-length sequences and represents each tile as a hash digest of the sequence it contains (Grishin et al. 2018). Data storage and access control is implemented using Blockstack. Blockstack makes use of blockchain, but not for storage; files are stored in a local drive or in the cloud (Digital Ocean, S3, Dropbox). The tile library of individual genomes (referenced by hash arrays) is stored in public storage such as InterPlanetary File System. Ethereum Smart Contracts are used to communicate among nodes, survey participants, and purchase access to data, but not to store data. (Grishin et al. 2018; Blockstack docs; Defrancesco and Klevecz 2019). As in the case with Zenome, Nebula uses blockchain to keep track of transactions and provide incentive for individuals to share their data. On the other hand, SAMchain uses blockchain to store raw genomics data on-chain.

**Cancer Gene Trust**: The Cancer Gene Trust (CGT) was built in 2019 (Defrancesco and Klevecz 2019), and is currently being run as a pilot program through UCSF. Their documentation describes an "a lightweight, global off-blockchain decentralized network controlled by "stewards" that make limited somatic mutation and related clinical data about a patient publicly available.” (Currie 2018). The data is stored off-chain in InterPlanetary File System, a peer-to-peer distributed file sharing system. Raw sequencing data is stored locally by data stewards. The blockchain component of CGT is Ethereum–a smart contract stores references to data files in the form a multihash of their content. Importantly, genomic data itself is not stored in Ethereum (references to the data are), and raw genomic data is not made available to the public CGT. Rather, blockchain is used as a tool to facilitate access to distributed files similar to CryptDist.(Cancer Gene Trust 2018; Currie 2018; Defrancesco and Klevecz 2019; Ozercan et al. 2018). SAMchain differs from CGT in that SAMchain stores NGS data on-chain to maximize data integrity.

**Encrypgen/Gene-Chain**: Encrypgen is a company founded in 2016 (Picco 2019) which has released Gene-Chain. Gene-Chain is a self-described DNA data marketplace. Their primary market is current users of 23andMe (Gonzalez and Kopsell 2020). Gene-Chain allows data owners to receive monetary compensation for access to their data via DNA tokens, though exactly how this works is not fully explained (Ozercan et al. 2018). From the available code, we can tell that the blockchain platform used is MultiChain. However, the code published here appears to be the multichain-web-demo repository (published by multichain), rather than the code showing how Gene-Chain works. Their use of MultiChain is also confirmed in their white paper (EncrypGen 2018). However, this white paper does not include any description of technical specifications. It is not clear whether the data will be stored in a MultiChain stream, or elsewhere. (Ozercan et al. 2018; encrypgen 2017; EncrypGen 2018; Defrancesco and Klevecz 2019). Due to the limited information available, we are not able to make a fair comparison between our application and Gene-Chain. However, we can tell that one key difference is the focus of the application. While Gene-Chain focuses on compensation and transactions, SAMchain focuses on high-integrity data storage and efficient querying.

In summary, most of the existing biomedical blockchain applications use blockchain for its cryptocurrency & transaction properties, while SAMchain and SCtools utilize it for data integrity purposes. In Table 3, we provide a concise comparison of these four blockchain platforms along with SAMchain.

### 1.2 Storing metadata vs. the data itself on the blockchain:

Here we present a simple analogy to illustrate why storing metadata and links to genomic data does not achieve data security. We can think of genomic data as books in a library. We can log the location (i.e. aisle) of each book in the library using a secure software. However, the books themselves will still be physically kept in the library, a single location. If there were a fire in the library, the logged information would not be affected, but we would still lose all of the books. In the same way, if we kept only the links to genomic data in a blockchain and the data itself elsewhere in a centralized storage system, the data would be lost if the centralized storage system were to fail. In other words, the security properties of blockchain would not be beneficial to the data itself.

### 2.1 MultiChain storage in a multi-node network

On a given node, MultiChain keeps a “wallet” directory on the disk, in addition to the blocks, which stores transactions of particular relevance to that node (i.e. transactions initiated by that node). This protocol is inherited from Bitcoin, however Bitcoin holds these transactions in memory while MultiChain stores them on disk (MultiChain 2020: Announcing the new MultiChain wallet). For example, let us say that a MultiChain network consists of three nodes A, B, and C, each node syncing the chain. If any of the three nodes publishes data to a stream, that data will be embedded in a block stored on each node. However, if node A is the one to publish all the data, node A will use more storage than B and C because in addition to storing the blocks, node A will also keep these transactions in its wallet. In the SAMchain network ecosystem, the vast majority of transactions happen at the initialization of the chain when the SAM data is published by the sequencer node. Thus, in Figure 2 Panel b, the first node (which published the SAM data to the chain) requires ~23 GB for this particular test dataset, while nodes 2-4 each require ~7.5 GB to sync the blocks.
